# Plant-microbe specificity varies as a function of elevation

**DOI:** 10.1101/540799

**Authors:** Gerald M. Cobian, Cameron Egan, Anthony S. Amend

**Author notes:** Correspondence: Anthony S Amend, Department of Botany, University of Hawaii Manoa, 3190 Maile Way, Honolulu, HI 96822 U.S.A.

## Abstract

Specialized associations between interacting species are a fundamental determinant of the diversity and distribution of both partners. How specialization of *guilds* of organisms varies along environmental gradients underpins popular theories of biogeography and macroecology, whereas the degree of specialization of a *species* is typically considered fixed. However, the extent to which environmental context impacts specialization dynamics is seldom examined empirically. In this study, we examine how specialization within a bipartite network consisting of three co-occurring plant species and their foliar fungal endophyte symbionts changes along a 1 000-meter elevation gradient where host species were held constant. The gradient, along the slope of Mauna Loa shield volcano, represents the entire elevational range of two of the three plants. Network and plant specialization values displayed a parabolic relationship with elevation, and were highest at middle elevations, whereas bipartite associations were least specific at low and high elevations. Shannon’s diversity of fungal endophytes negatively correlated with specificity, and was highest at the ends of the transects. Although plant host was a strong determinant of fungal community composition within sites, fungal species turnover was high among sites and plant host predicted a weak, though significant proportion of compositional variance. There was no evidence of spatial or elevational patterning in fungal community compositon. Our work demonstrates that specificity can be a plastic trait which is influenced by the environment and centrality of the host within its natural range.

## Introduction

Plant microbe symbioses vary in the degree to which one or more partners “specializes” on the other. Some hosts only associate with specific symbiont species such as mycoheterotrophic plant species from the subfamily Monotropoideae where each associates with a different, but specific, ectomycorrhizal fungal partner [1]. Other hosts are specific at higher taxonomic ranks such as the legume-rhizobia symbiosis [2] or species of Pinaceae that are the only hosts of the ectomycorrhizal fungal genus *Rhizopogon* [3]. Other plant hosts and fungal symbionts form more general associations, such as most ectomycorrhizal fungi which form symbioses with many woody host plants [4] and arbuscular mycorrhizal fungi, where fewer than 300 species associate with ~70% of land plant families [5][6]. Foliar pathogens, as a group, lie somewhere in between, as many species encompass the ability to infect a broad diversity of hosts in some circumstances, but the probability of infecting two host plants diminishes as a function of phylogenetic distance [7].

One method in which the degree of specialization between a host and a microbial symbiont can be quantified is by accounting for the frequency of the interaction between two species relative to other community members. In nature these species interactions act on a continuum ranging from complete generalization to full specialization [8]. Placement along this continuum presents a series of tradeoffs. For example, in the case of mutualisms, if symbionts are optimally suited for each other, collocated, and abundant, specialization can confer a fitness benefit since energy is not wasted maintaining associations with inferior partners. However, if one symbiotic partner is sparse or absent, specialization comes at a cost, since it might limit the distribution or fitness of the other partner.

The degree to which specialization varies across systems and environments has long been considered an important component of latitudinal diversity clines [9, 10]. In fact, analyses of interactions across broad geographic scales provide some evidence for this idea [11, 12]. Broad-scale measures of species interactions, however, are minimally informative for interactions happening at the local scale, and there is little evidence for how environmental gradients impact specificity within a given species or community [13]. This gap is likely attributable to the fact that species composition typically covaries as a function of environment such that specialization as a function of the environment cannot be disentangled from specialization as a function of species identity.

Elevation gradients are an ideal system in which to examine how specialization between plant hosts and microbial symbionts covaries with changes in environmental conditions, as the same host species can be found along the entirety of a gradient along with coinciding large variations in abiotic conditions [14]. In this study, we took advantage of a steep elevation gradient on the Island of Hawai□i to examine how foliar fungal endophyte host specialization is influenced by environmental conditions. Specifically, we examined foliar fungal endophyte hosts along Mauna Loa, a shield volcano on Hawai□i Island. The volcano forms a steep environmental gradient as it rises from sea level to 4 200 meters above sea level (masl) over approximately 20 km. Additionally, the young geological age and isolation of the Hawaiian archipelago supports a relatively species poor (circa 1 100 spp.) but primarily endemic flora, some of which encompass wide geographical niches and elevational distributions [15]. For example, *Metrosideros polymorpha*, an ecologically important and endemic tree species, can be found near sea level to 2 500 m along the eastern slope of Mauna Loa [16]. In addition to *M. polymorpha*, there are several other woody species that co-occur along the gradient enabling us to examine wide environmental variance while keeping host identity constant.

The foliar fungal endophyte symbiosis, defined here as all fungi living *within* leaf tissue but not causing any outward signs of disease [17], are effectively invisible but represent one of the most ubiquitous symbioses in nature. Researchers have yet to find a plant lineage lacking these cryptic microbial associates. Endophytes can play important roles in plant biochemistry (reviewed: ref. 17), water conductance [19], heat and drought tolerance [20], and disease resistance [21]. Despite a presumed horizontal transmittance [22] foliar fungal endophytes demonstrate varied degrees of specialization among plant hosts. In one study spanning North America [23] fungal community dissimilarity correlated with phylogenetic dissimilarity of hosts, and in another, plant species in lowland tropical forests in New Guinea [24] showed more similar fungal community composition within-host species compared to among. Within controlled experiments, foliar fungal endophyte composition can vary among host plants at the population level [25, 26]. However, the extent to which the environment affects the strength of specialization in this host-microbe symbiosis is largely unknown.

We combined the unique characteristics of the Hawaiian flora with the steep elevation gradient of Mauna Loa to examine how changes in environmental conditions can affect host specialization, local richness, and community composition. Because environmental conditions along Mauna Loa become more stressful with increasing elevation (e.g. decreased precipitation, increased solar radiation, and greater diurnal temperature differences), we expected to observe a decrease in fungal species richness with increasing elevation, as fewer species would be able to tolerate and persist under conditions at higher elevations. Based on previous studies examining host-associated microbes, we hypothesized that elevational gradients would structure foliar fungal endophyte community composition, such that communities located at more similar elevations will be more similar in composition. Furthermore, we predicted that host identity would play an important role within and among sites in determining fungal community composition. Finally, we predicted that specificity (both network and individual hosts) would decrease as a linear function of elevation, which is consistent with observations of mutualistic interaction network specialization patterns along latitudinal gradients [27].

## Materials and Methods

### Sites/Fieldwork

Sampling was conducted along the Eastern slope of Mauna Loa on Hawai□i Island (19.4721° N, 155.5922° W; Figure 1). Along this steep slope the volcano encompasses an environmental gradient over which temperature, rainfall and solar irradiance vary rapidly (Figure S1). To examine the influence of the abiotic environment on foliar endophyte host specialization we sampled three native Hawaiian plants that co-occur along the gradient. Hosts were sampled from 1 100-2 000 masl at 100 m intervals and included: *Leptecophylla tameiameiae* (pūkiawe) an indigenous species found throughout the Pacific, *Metrosideros polymorpha* (□ōhi□a) an endemic Hawaiian species common throughout the Hawaiian archipelago, and *Vaccinium reticulatum* (□ōhelo) also a Hawaiian endemic. For both *L. tameiameiae* and *V. reticulatum*, our sampling range encompassed the entire extent of their elevation distribution along this portion of Mauna Loa. *M. polymorpha* is the dominant tree species along the entire gradient, and its range extends both above and below the limits of sampling conducted for this study. Within each of the ten plots sampled, we collected leaf samples from four different individuals of each host species (120 sampled individuals, 40 per host species total).

**Figure 1.**
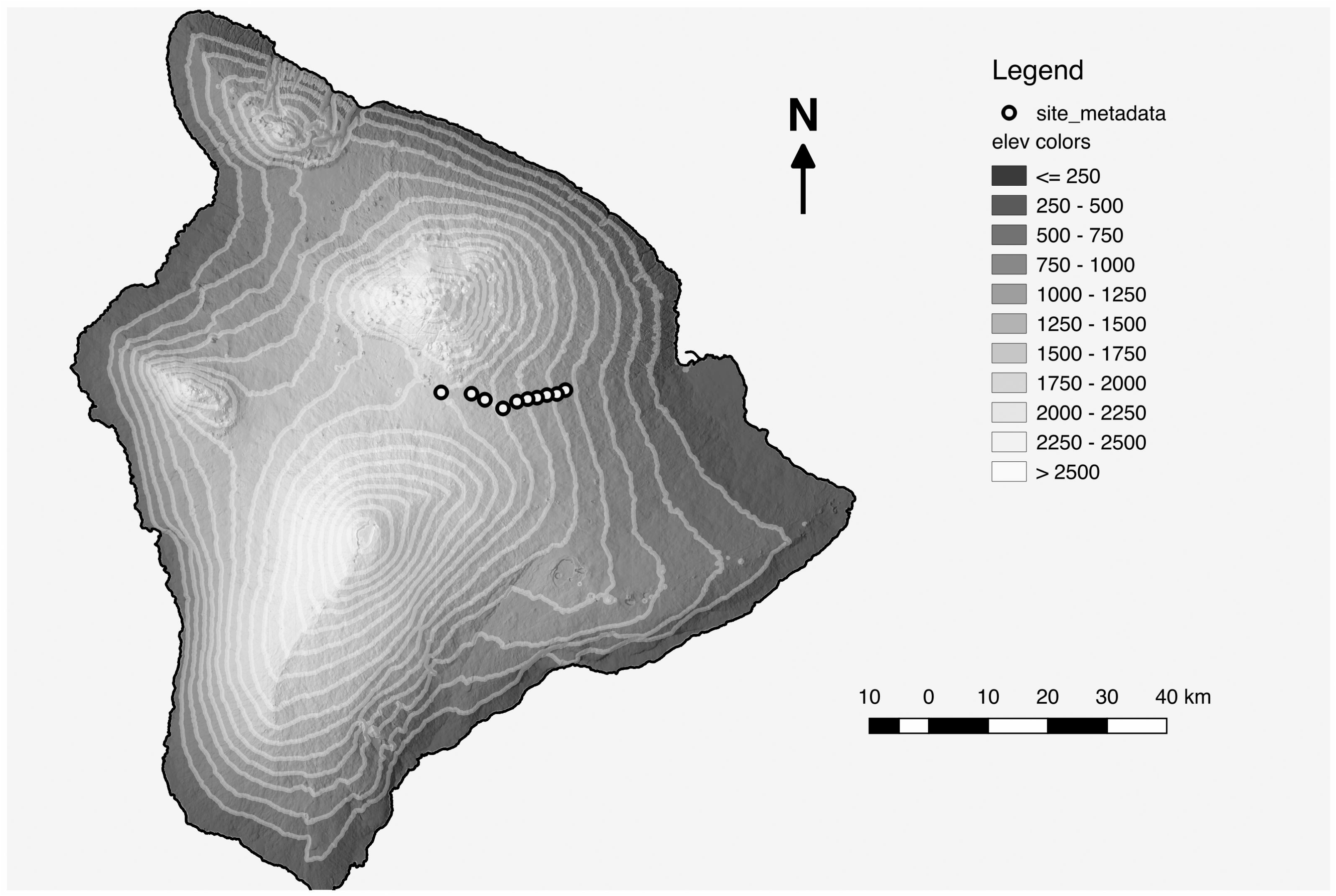
Sampling locations on the Island of Hawai□i. Circle points indicate sampling locations, shades of grey indicate elevation in meters above sea level (masl). Starting at 1 100 masl, sampling locations were established in 100 m increments spanning approximately 20 km to our highest elevation site at 2 000 masl. At each site, we collected leaf samples from four individuals for each of our target host (*L. tameiameiae, M. polymorpha*, and *V. reticulatum*).

Sampling was based on approximate leaf mass. We haphazardly collected 5 apparently healthy leaves from *M. polymorpha* and *V. reticulatum* individuals, and 40 leaves from *L.* tameiameiae individuals (as its leaves are much smaller). Leaf samples were placed in Ziploc bags and stored on ice in the field, before being placed in −20 °C storage in the lab. Within each elevation, we collected a voucher specimen from each host species and deposited them in the Joseph F. Rock Herbarium at the University of Hawai□i at Mānoa.

### Molecular analysis

#### Surface sterilization

For each replicate we collected forty leaf-disks per individual by punching leaves using a surface sterilized standard paper single hole punch (approximately 0.5 cm diameter). Because *L. tameiameiae* leaves are roughly 0.7 cm long, entire leaves of this species were processed. Leaf disks and *L. tameiameiae* leaves were placed inside loose-leaf tea bags, and surface-sterilized by submerging in 1% NaClO for 2 mins, 70% EtOH for 2 mins, and finally two rinses in sterile water for 2 mins each. Rinse water was retained for use as a negative control in subsequent analyses.

#### DNA Extraction

For DNA extraction, ten surface sterilized leaf-disks per sample were haphazardly chosen from tea bags and placed in MP Biomedical Lysing Matrix A tubes (MP Biomedical, Santa Ana, CA, USA) containing Solution PD1, Solution PD2, Phenolic Separation Solution, and RNase A Solution from the MoBio PowerPlant Pro DNA Isolation kit (MO Bio, Carlsbad, CA, USA). Leaf disks were homogenized using a Mini-Beadbeater 24 (BioSpecs Inc. OK, USA) at 3 000 oscillations per min for 2 mins. Genomic DNA was then isolated using the MoBio PowerPlant Pro DNA Isolation kit protocol following the manufacturer’s instructions.

#### Amplification and Illumina Library Prep

We amplified the ITS1 region of the ribosomal RNA gene (rDNA) using the fungal specific primers ITS1f and ITS2, along with Illumina adaptors and Golay barcodes following the protocols of Smith and Peay [28], but with the addition of a second 12 bp nucleotide index on the ITS1f primer construct to enable a dual indexed approach. Reaction conditions were as follows: initial denaturation at 95 °C for 180 sec followed by 35 cycles of 95 °C for 20 sec, 53 °C for 15 sec, and 72 °C for 20 sec, with a final elongation at 72 °C for 60 sec. PCRs were carried out in 25 *µ*L PCR reactions containing 1X KAPA3G Plant PCR kit master mix (KAPA Biosystems, Wilmington, MA, USA), 1.5 mg/mL BSA, 2 mM MgCl2, 0.3 *µ*M forward primer, 0.3 *µ*M reverse primer, 0.2 *µ*L KAPA3G enzyme, and 9 *µ*L DNA template. PCR amplification was verified via gel electrophoresis. PCR products were purified and normalized using Just-a-Plate^™^ 96 PCR Purification and Normalization Kit (Charm Biotech, San Diego, California, USA) and concentrated using a streptavidin magnetic bead solution. The resulting library was sequenced by GENEWIZ (GENEWIZ, South Plainfield, NJ, USA) using the 2 x 300 paired-end (PE) sequencing chemistry on an Illumina MiSeq sequencing platform (Illumina Inc., San Diego, CA, USA). Raw data have been deposited into the National Center for Biotechnology Information’s Sequence Read Archive (PRJNA474551).

### Bioinformatics

Bioinformatic analyses were conducted using Quantitative Insights Into Microbial Ecology (QIIME v1.9; ref. 28) and Mothur v1.39 [30]. Sequencing produced 23 531 047 sequences. After removal of sequence with an average Quality score <25 or a sequence length <75, 18 370 578 were retained. Because reverse reads contained a high proportion of low-quality reads, only forward reads were utilized for subsequent processing and analyses.

Forward sequences were clustered into operational taxonomic units (OTUs) using a chain-clustering protocol. We chose this method as it tends to be more accurate compared with using a single OTU clustering algorithm [31]. Sequences were first grouped into *de novo* OTUs using a sequence similarity of ≥96% using USEARCH [32]. Chimera checking referenced the UNITE v7 database [33]. A second round of *de novo* clustering was then performed using UCLUST [32], again using ≥96% sequence similarity. Representative sequences from OTUs were then assigned taxonomy using the BLAST v2.6 algorithm [34] and the UNITE v7 database within QIIME. Query sequences aligning to less than 85% of the length of a fungal database sequence were removed from the OTU table. No OTUs were detected in PCR negative controls after OTU clustering. We then exported our OTU table to R (v3.3.2; ref. 34) for further processing.

To reduce the likelihood of mis-attribution due to tag switching during the sequencing run, we removed OTUs from samples in which their abundance was <0.1% of the maximum number of reads found in another sample. We then filtered all OTUs with <10 reads in the dataset, as well as OTUs not identified to the kingdom of Fungi. This resulted in 2 058 OTUs. We then down-sampled our OTU table to 2 300 reads per sample to account for uneven sequencing depth across the dataset. While there is no consensus on how best to account for differences in sequencing depth across samples, given a mean per sample richness of fewer than 60 OTUs in our study, we believe that down-sampling to achieve equal sequencing coverage minimally impacts statistical sensitivity.

### Statistical Analyses

#### Effect of elevation on local fungal richness and Shannon diversity (alpha-diversity)

We examined local alpha-diversity of foliar fungal endophytes using both observed species richness and Shannon’s diversity index as our metrics. Both diversity metrics were calculated within individual plants. Species richness was calculated using the *specnumber* function and Shannon diversity was calculated using *index = “shannon”* in the *diversity* function, both of which are available in the vegan package [36]. Changes in alpha-diversity with increasing elevation were examined along our gradient using Pearson’s moment correlation with the *cor.test* function in base R [36].

#### Effects of elevation and host identity on fungal community composition (beta-diversity)

To examine differences in community composition among samples we first calculated Bray-Curtis [37] dissimilarity among samples using log-transformed OTU abundances. To determine the changes in fungal community composition along our gradient, we calculated Mantel correlations [38] between elevational and community compositional dissimilarities using the *mantel* function with 10 000 permutation in the vegan package [36]. To further quantify changes in beta-diversity along our gradient, we determined the relative contribution of species nestedness vs species turnover separately for all three hosts [39]. Changes in beta-diversity were partitioned into the nestedness and turnover components using the *beta.multi.abund* function in the betapart package [40].

To determine the extent to which host identity predicts community composition within each site along the gradient we performed a permutational multivariate analysis of variance (PERMANOVA; ref. 40) using the *adonis* function in the Vegan package [36].

#### Host specialization and selectivity

To determine how host specialization of foliar fungal endophytes changes as a function of elevation, we implemented a network approach. We assembled bipartite networks by aggregating fungal OTUs by host species within each elevation. We calculated the H2’, a measure of specialization [8] generalized across the entire network, for each elevation. The index describes the extent to which observed interactions deviate from those that would be expected given the species and sample abundance distributions of species [8]. H2’ values range from 0 (no network specialization) to 1 (perfect network specialization) based on potential associations given OTU abundance totals. H2’ was calculated using the *H2fun* function in the *bipartite* package [42]. Next, we examined plant host specialization for foliar fungal endophytes for our three targets hosts separately. Host specialization was determined using the d’ specialization index using the *dfun* function [8]. The d’ index calculates how strongly a species deviates from a random sampling of interacting partners available, and like H2’ it ranges from 0 (no specialization) to 1 (perfect specialist) [8]. Both indices consider interaction frequency (abundance) and are standardized to account for heterogeneity in the interaction strength and taxon richness. To examine the relationship between elevation and H2’ and d’, we modeled both indices with increasing elevation using quadratic polynomial regression. For each polynomial regression, model assumptions were verified using the *gvlma* function in the gvlma package [43].

To test whether observed networks exhibited non-random patterns we used a null model approach. To generate null networks, we randomized observed networks 10 000 times using the *swap.web* algorithm and the *nullmodel* function in the bipartite package and compared empirical values to the randomized distribution using a one-sample t-test.

## Results

### Effect of elevation on local fungal richness and Shannon diversity (alpha-diversity)

With the exception of endophytic communities associated with *L. tameiameiae*, which showed a weak, negative correlation between diversity and elevation (Figure 2, *r* = −0.325, *p* = 0.041) we observed no significant relationship between fungal species richness or diversity and elevation among sampled hosts (Figure 2).

**Figure 2.**
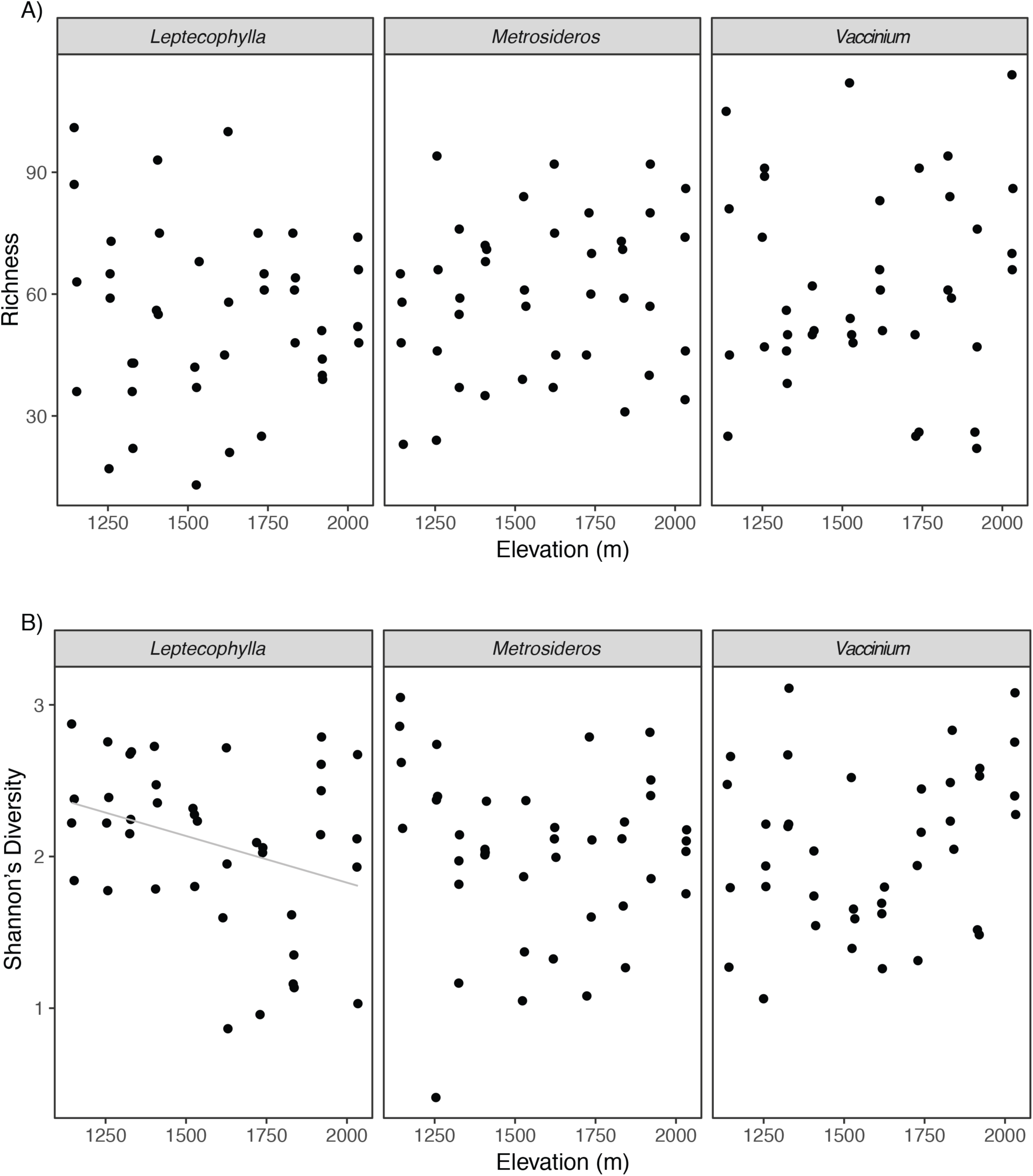
Top three panels (**A**) display fungal richness associated with our three target hosts plotted against elevation. Fungal richness was not affected by elevation (*L. tameiameiae*: R^2^ = 0.003, p = 0.731; *M. polymorpha*: R^2^ = 0.029, p = 0.295; *V. reticulatum*: R^2^ = 0.055, p = 0.738). Bottom three panels (**B**) display fungal community Shannon’s diversity associated with our three target hosts plotted against elevation. Fungal Shannon’s diversity for both *M. polymorpha* and *V. reticulatum* were not significantly influenced by elevation (R^2^ = 0.01, p = 0.545; R^2^ = 0.052, p = 0.160; respectively). Shannon’s diversity of fungal communities associated with *L. tameiameiae* declined significantly with increasing elevation (R^2^ = 0.105, p = 0.041). Solid gray line represents linear regression.

### Effects of elevation and host identity on fungal community composition (beta-diversity)

Overall, elevational dissimilarity was a poor predictor of endophyte compositional similarity (Figure S2; Mantel test for correlation between two distance matrices; *r* = 0.023; *p* = 0.119; Permutations = 10 000). Neither *M. polymorpha* (Figure S2; Mantel test for correlation between two distance matrices; *r* = 0.099; *p* = 0.054; Permutations = 10 000), nor *V. reticulatum* (Mantel test for correlation between two distance matrices; *r* = 0.089; *p* = 0.056; Permutations = 10 000) associated communities demonstrated a significant relationship between elevational dissimilarity and foliar endophytic fungal community composition although communities associating with *L. tameiameiae* were weakly correlated (Figure S2 Mantel test for correlation between two distance matrices; *r* = 0.105; *p* = 0.018; Permutations = 10 000).

In contrast, host identity strongly determined foliar fungal endophytic community composition at all elevations sampled, with the exception of 1 700 and 2 000 masl (Figure 3A, Table 1). Host identity explained the greatest amount of variance at mid elevations (1 300 m – 1 600 m), and then decreased at lower and higher elevations (Table 1). When all elevations were considered, host identity was a significant though weak predictor of fungal community composition (Table S1; Figure S3; ADONIS: *R*^*2*^ = 0.104, *p* < 0.001). When we examined the relative contribution of species turnover vs nestedness in determining changes in community composition, we observed that changes along our gradient were driven primarily by species turnover for all hosts (Table 2).

**Table 1.** Variation in foliar fungal endophyte community composition (beta (β) diversity) among hosts within each elevation sampled, as determined by permutational multivariate analysis of variance using distance matrices (PERMANOVA). Values shown include degrees of freedom (DF), sum of squares (SS), and mean squares (MS). Asterisks indicate significance at p < 0.05.

| **Elevation**<br>**(masl)** | **β-diversity**<br>**Bray-Curtis** | **DF** | **SS** | **MS** | **PseudoF/F** | **R2** | **p value** |
| --- | --- | --- | --- | --- | --- | --- | --- |
| **1100** | Host | 2 | 1.155 | 0.578 | 1.849 | 0.291 | 0.006 ** |
|  | Residuals | 9 | 2.813 | 0.313 |  | 0.709 |  |
| **1200** | Host | 2 | 1.119 | 0.375 | 1.493 | 0.249 | 0.033 * |
|  | Residuals | 9 | 3.372 |  |  | 0.751 |  |
| **1300** | Host | 2 | 1.518 | 0.759 | 2.085 | 0.317 | 0.002 ** |
|  | Residuals | 9 | 3.277 | 0.364 |  | 0.683 |  |
| **1400** | Host | 2 | 1.858 | 0.929 | 3.642 | 0.477 | 0.001 *** |
|  | Residuals | 9 | 2.040 | 0.255 |  | 0.523 |  |
| **1500** | Host | 2 | 2.136 | 1.068 | 3.751 | 0.455 | 9.999E-05 *** |
|  | Residuals | 9 | 2.563 | 0.285 |  | 0.545 |  |
| **1600** | Host | 2 | 1.779 | 0.890 | 3.084 | 0.407 | 0.002 ** |
|  | Residuals | 9 | 2.595 | 0.288 |  | 0.593 |  |
| **1700** | Host | 2 | 0.710 | 0.355 | 0.850 | 0.159 | 0.623 |
|  | Residuals | 9 | 3.756 | 0.417 |  | 0.841 |  |
| **1800** | Host | 2 | 1.408 | 0.704 | 2.283 | 0.337 | 0.013 * |
|  | Residuals | 9 | 2.776 | 0.308 |  | 0.663 |  |
| **1900** | Host | 2 | 1.230 | 0.615 | 1.658 | 0.269 | 0.019 * |
|  | Residuals | 9 | 3.339 | 0.371 |  | 0.731 |  |
| **2000** | Host | 2 | 0.734 | 0.367 | 0.949 | 0.174 | 0.520 |
|  | Residuals | 9 | 3.480 | 0.387 |  | 0.826 |  |

**Table 2.** Partitioned abundance-based beta (β) diversity metrics for foliar fungal endophyte community composition within each host examined along the gradient. β_BC-bal_ indicates changes in community composition caused by balanced variation in abundance, whereby the individuals of some species in one site are substituted by the same number of individuals of different species on another site (i.e. species turnover) [39]. β_BC-gra_ indicates the changes in community composition caused by abundance gradients, whereby some individuals are lost from one site to the other (i.e. community nestedness) [39]. β_BC_ indicates changes in community composition as determined by Bray-Curtis dissimilarity [37].

| Measure of compositional<br>dissimilarity (beta diversity) | Host |  |  |
| --- | --- | --- | --- |
|  | *L. taeiameiae* | *M. polymorpha* | *V. reticulatum* |
| Turnover ($\beta_{\text{BC-bal}}$) | 0.975 | 0.960 | 0.965 |
| Nestedness ($\beta_{\text{BC-gra}}$) | 0.000 | 0.000 | 0.000 |
| Overall ($\beta_{\text{BC}}$) | 0.975 | 0.960 | 0.965 |

**Figure 3.**
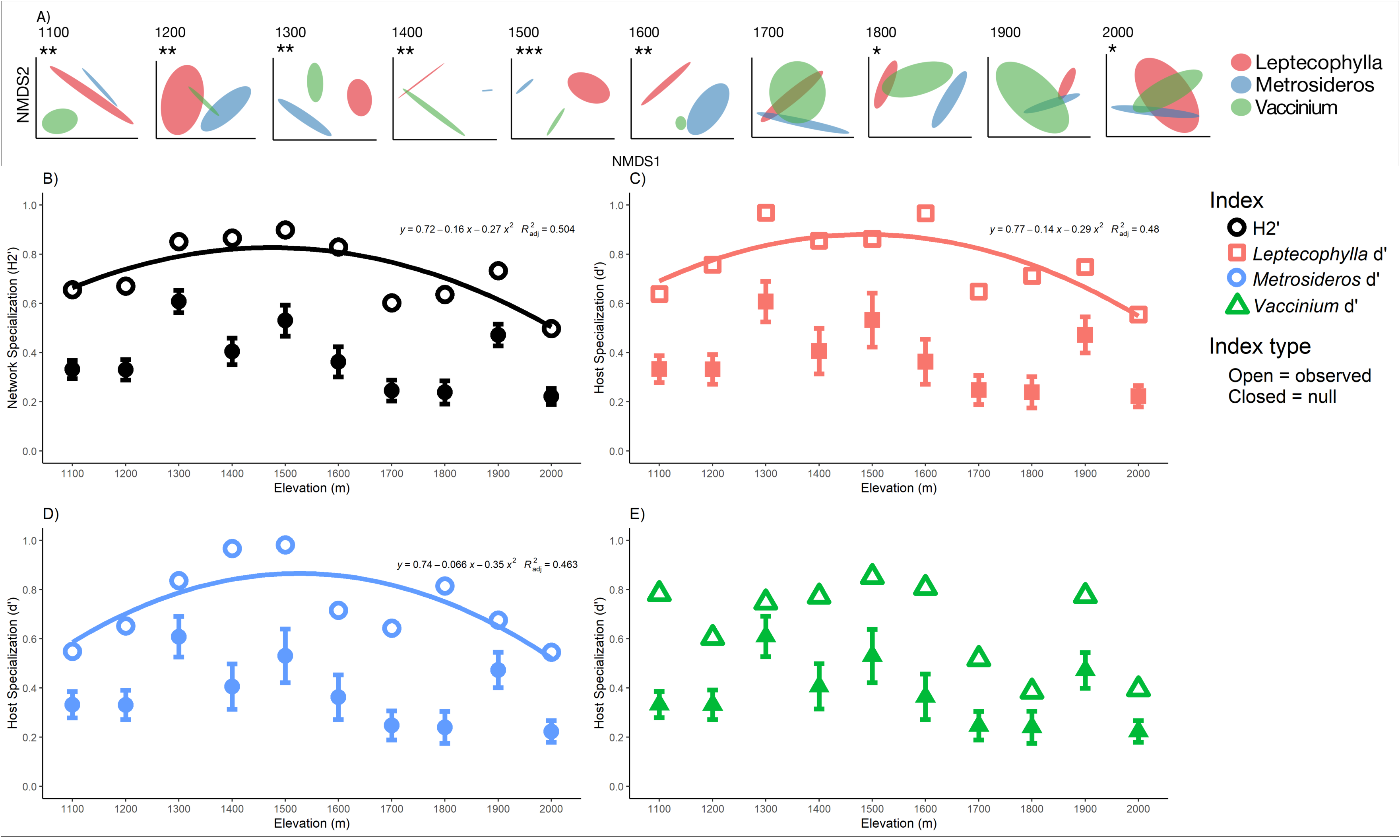
Top panel (**A**) displays NMDS plots for each elevation site and illustrates how endophyte communities are structured by host identity. Shaded ellipses represent standard error of the mean (95%) and are colored by host (red = *L. tameiameiae*, blue = *M. polymorpha*, green = *V. retaculatum*). Asterisks above NMDS plots show significance values of PERMANOVA tests, where: * ≤0.05, ** ≤0.01, *** ≤0.001). Lower four panels (**B, C, D, E**) display observed; network specialization (H2’; **B**, open black circles), and host specialization (d’) for *L. tameiameiae* (**C**, open red squares), *M. polymorpha* (**D**, open blue circles), and *V. retaculatum* (**E**, open green triangles) respectively plotted against elevation. For all network indices, significant trends (at p < 0.05) with elevation are illustrated with a polynomial regression curve fitted to the graph (polynomial order = 2), along with associated quadratic equation and adjusted-R^2^ value. Observed network indices were compared to random expectation by assembling null networks using the *swap.web* algorithm and the *nullmodel* function with 10 000 permutations in the bipartite package (Dorman et al. 2008). Null means for each network index at each site are denoted by closed shapes, along with standard deviations around the null mean (error bars around closed shapes). Observed H2’ and d’ for all three species were significantly more specialized than expected by chance at all sites (see Table S2 and S3 for t-test results).

### Host specialization and selectivity

Network specialization displayed a unimodal pattern along the elevation gradient sampled (Figure 3B, Table S2), where highest network specialization values were observed at mid elevations, and then decreased at higher and lower elevations. Network specialization peaked at 0.9 at 1 500 masl and then was lowest at 0.5 at 2 000 masl (Figure 3B, Table S2). Network specialization could be well fitted to a polynomial regression equation (Figure 3B; Polynomial order = 2; *R*^*2*^_*adj*_ = 0.0.504; F-statistic = 5.572; df = 2,7; *p* = 0.036). Indicating that elevation is a strong predictor of network specialization between plants and foliar fungal endophytes.

Like network specialization, host specialization on foliar fungal endophytes for all three hosts displayed a unimodal pattern with elevation, where specialization for all three species peaked between 1 400 and 1 500 masl and decreased at lower and higher elevations (Table S3). The lowest host specialization for all three-host species was observed at the highest elevation sampled (Table S3). Both *L. tameiameiae* (Figure 3C; red curve; Polynomial order = 2; *R*^*2*^_*adj*_ = 0.0.4802; F-statistic = 5.157; df = 2,7; *p* = 0.040) and *M. polymorpha* (Figure 3D; blue curve; Polynomial order = 2; *R*^*2*^_*adj*_ = 0.4628; F-statistic = 4.877; df = 2,7; *p* = 0.047) specialization fit a polynomial regression equation. In contrast, *V. reticulatum* was poorly fitted by a polynomial regression equation (Figure 3E; Polynomial order = 2; *R*^*2*^_*adj*_ = 0.174; F-statistic = 0.1732; df = 2,7; *p* = 0.21), indicating that elevation strongly influenced *L. tameiameiae* and *M. polymorpha* specialization on fungal foliar endophytes, but not *V. reticulatum*. Null model analysis revealed that within all sites along our gradient observed network specialization (Figure 3B, Table S2), as well as observed specialization for all hosts (Figure 3C, 3D, 3E, Table S3) were significantly higher than expected by chance (P<0.001).

## Discussion

Host specificity is typically considered a static trait, although environmental impacts on biotic interaction patterns and strength are seldom evaluated in experimental systems. Here, we provide evidence for elevational clines in specialization. We observed that both total network specialization (H2’), as well as individual host specializations (d’) peaked at mid-elevations and then decreased towards the ends of the elevation gradient we examined. This pattern ran contrary to our prediction that specificity would decrease linearly as a function of elevation, which is consistent with patterns found for other species interactions along latitudinal gradients [27]. Although future empirical work will be needed to examine the mechanisms of host specificity along elevational gradients, we consider two plausible and non-exclusive mechanisms that might explain the pattern observed.

First, we hypothesize that fungal endophyte specialization is influenced by host species density. Host density-dependent disease dynamics predict a higher occurrence of host specific disease when hosts are more abundant, and specificity of disease decreases as host species become more scarce [10, 44]. Because foliar fungal endophytes employ similar mechanisms to associate with hosts as fungal pathogens, mainly unrestricted dispersal of gametes and sexual reproduction before infection of a new host [45], we expect they are highly likely to be affected by host density. Host species occurrences are patchier at the edges of their ranges, whereas at mid elevations, they are more regularly distributed and abundant. Many fungi exhibit a tradeoff between dispersal and competitive ability[46]. Assuming that specialized interactions confer a competitive advantage, the higher host specialization found at mid elevations might be due to decreased reliance on dispersal ability exhibited by more generalist fungi at low and high elevations.

Additionally, fungal endophyte communities may be less specialized at lower and higher elevations because these represent the limits of the host species range. Populations at the margins of their species’ distributions tend to be at lower abundances than populations in the middle of their species range distributions [47, 48]. As such, marginal populations also tend to have elevated physiological stress and lower genetic diversity [49] compared to populations near the middle of their range distribution, which in turn leads to lower population growth rates and higher mortality [50]. Because our gradient constituted the entirety of the elevational range of two of the host species examined (and the majority of the third), we suspect that sampled plants experience increased stress at either end of the elevation gradient. Although physiological mechanisms governing host control of microbial symbionts, particularly leaf symbionts, is poorly understood, plant innate immunological responses are metabolically expensive and require the (re)allocation of resources from other plant functions such as growth, reproduction, and stress responses [51]. Thus, hosts would likely have a better ability to allocate resources to select for and against symbionts when they are central in their ranges, whereas plants located at the margins will have fewer resources to be selective.

Although there are likely multiple mechanisms that reinforce endophyte specialization, numerous studies have indicated some degree of host control. Because endophytic fungi are contained within host leaves, they are in intimate contact with their hosts and receive continual chemical feedback. For example, endophytic fungi grow faster on media containing host leaf extract than non-host leaf extract [22, 52]. Our observance of hosts being a strong determinant of endophyte community across our gradient, combined with previous lab and greenhouse studies, suggest that hosts provide unique biotic environments that select for specific fungal partners, and that host selection remains a strong determinant of community composition regardless of local abiotic conditions.

In this study, we observed that host species significantly influenced fungal endophyte community composition both within and among elevations, contributing to the growing body of literature showing that endophytic communities are strongly selected for by host species [23, 24, 53, 54]. That host species are a stronger determinant of endophyte community composition than elevation was a surprising result, particularly when coupled with high species turnover across sites. That is to say, whereas host species was the primary determinant of fungal community composition within elevations, hosts associate with a largely unique set of fungal endophytes at each elevation that are dissimilar from both communities associated with co-occurring plant species, and from conspecific plants at other elevations.

In contrast, multiple studies show that local environmental conditions influence foliar fungal endophyte communities. For example, endophyte communities associated with two different grass species were structured by rainfall [55]. Similarly, Zimmerman and Vitousek [56] observed that foliar endophyte communities within *M. polymorpha* were structured by elevation and aspect (but not soil age) along a steep volcanic gradient on the Island of Hawai□i. Despite starkly different values of solar radiation, temperature, precipitation, humidity, and cloud cover at the ends of our gradients, we did not find evidence that endophyte communities were structured significantly by local abiotic conditions. One reason for this discrepancy may be geographic scale. Even though our study was conducted along a portion of the same gradient as Zimmerman and Vitousek, and with one of the same host species (*M. polymorpha*), we did not replicate their strong elevation partitioning. However, in their study Zimmerman and Vitousek observed that endophyte communities sampled from similar elevations and aspect (1 100 masl and 1 800 masl) were less differentiated compared to samples collected higher and lower along the transect. The niche breadth of *M. polymopha* is remarkably wide, and our sampling, limited by the elevational distributions of the other two plant species, did not capture the most divergent communities. Similarly, the large effect sizes noted by Giauque and Hawkes [55] considered hosts sampled across 400 km, nearly an order of magnitude greater than the distance sampled in this study. This increased scale may lead to greater effects as well an increased likelihood that both host phylogenetic variation and/or fungal dispersal limitation, in addition to environmental variance, might also contribute to their patterns noted. In fact, in a larger archipelago-wide survey of endophytes associated with >100 host plant species and spanning 2 300 m in elevation, we found that numerous environmental factors, principally elevation and plot level evapotranspiration, correlated with endophyte community composition (in revision).

Alpha diversity was similarly unaffected by elevation. For larger organisms, such as plants and animals, species richness tends to decrease with increasing elevation. However, correlations between elevation and diversity are less consistent among microbial communities. In this study, we observed no relationship between elevation and fungal endophyte species richness or diversity, congruent with other studies examining soil fungi [57], soil bacteria [58], or leaf bacterial communities [59], but in contrast to others [60, 61]. Collectively these results indicate that mechanisms structuring ‘micro’ organismal groups are not uniform, resulting in different diversity patterns along elevation gradients.

Foliar fungal endophytes form intimate associations with their plant hosts. While previous research has shown that both environment and host identity strongly influence endophyte community composition, we observed that the local abiotic environment has little influence on fungal richness or community composition compared with host species. This highlights the need for research spanning a diverse array of ecosystem types to generalize our knowledge of the factors structuring these ubiquitous plant associated microbes. We observed that host and network specialization, as well as community dissimilarity peak at mid elevations indicating that the strength of biotic interactions between plant and symbiont vary as a function of elevation. Collectively, our results show a strong interaction between host and endophyte, and that the environment influences fungal communities, indirectly, by modulating host specialization.

Supplementary information is available at ISME Journal’s website.

## Acknowledgements

The authors wish to thank Erin Datlof at UH Hilo and Tomoko Sakishima at UNLV for field assistance, and members of the Amend and Hynson labs for critical review of earlier versions of the manuscript. The authors also wish to thank the National Science Foundation for their generous support as NSF grant #1255972 to ASA and NSF grant #1329626 to GMC.

