## Supplementary material for "Plant-microbe specificity varies as a function of elevation": Figure S1

Supplemental Figure 1:

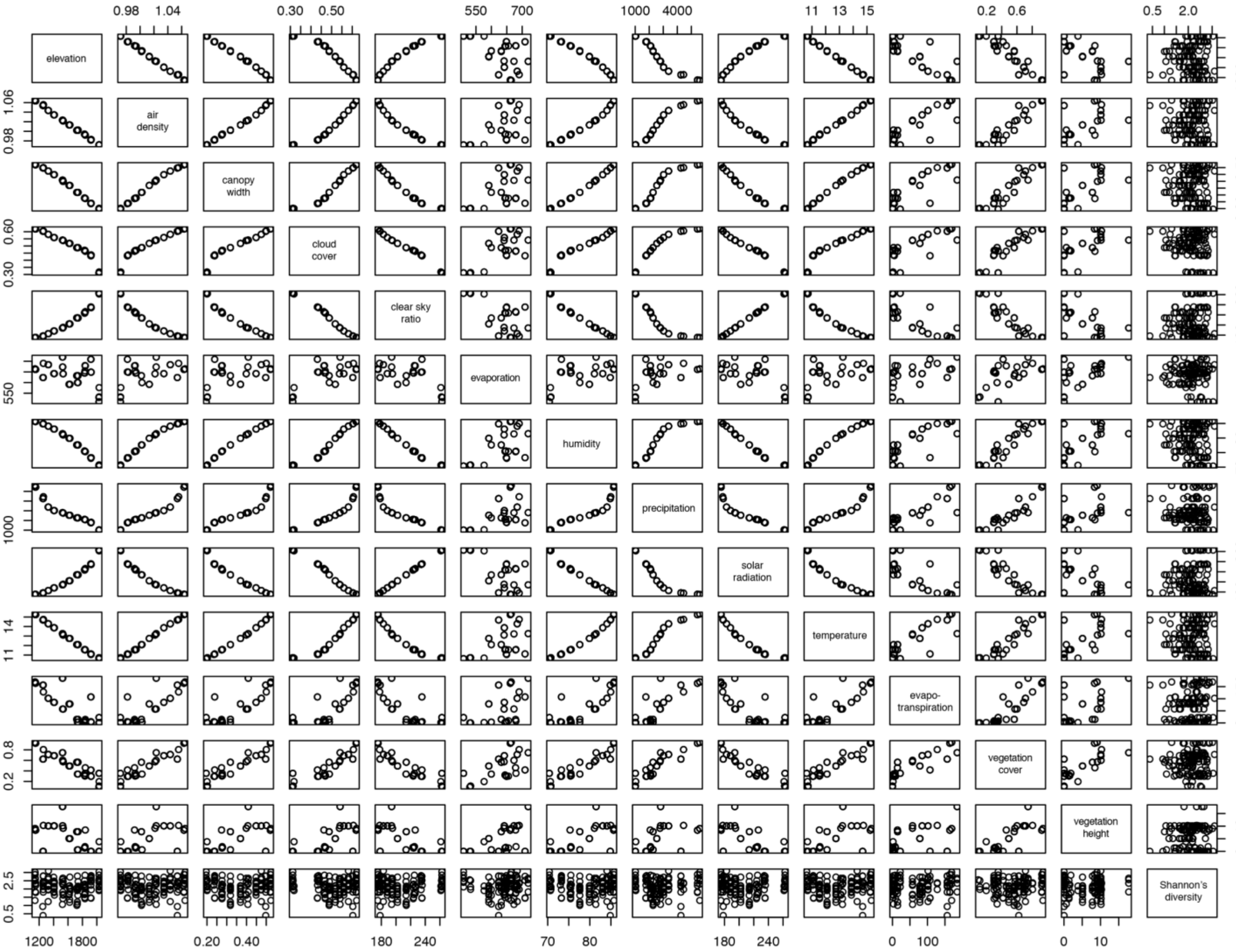


**Figure S1.** Matrix scatterplot shows the relationships of various environmental parameters of our elevation gradient. There were clear negative relationships between elevation and air density, canopy width, cloud cover, humidity, precipitation, temperature, evapotranspiration, and canopy cover. There were significant positive relationships between elevation and clear sky ratio and solar radiation.
