## Supplementary material for "Plant-microbe specificity varies as a function of elevation": Figure S2

Supplemental Figure 2:

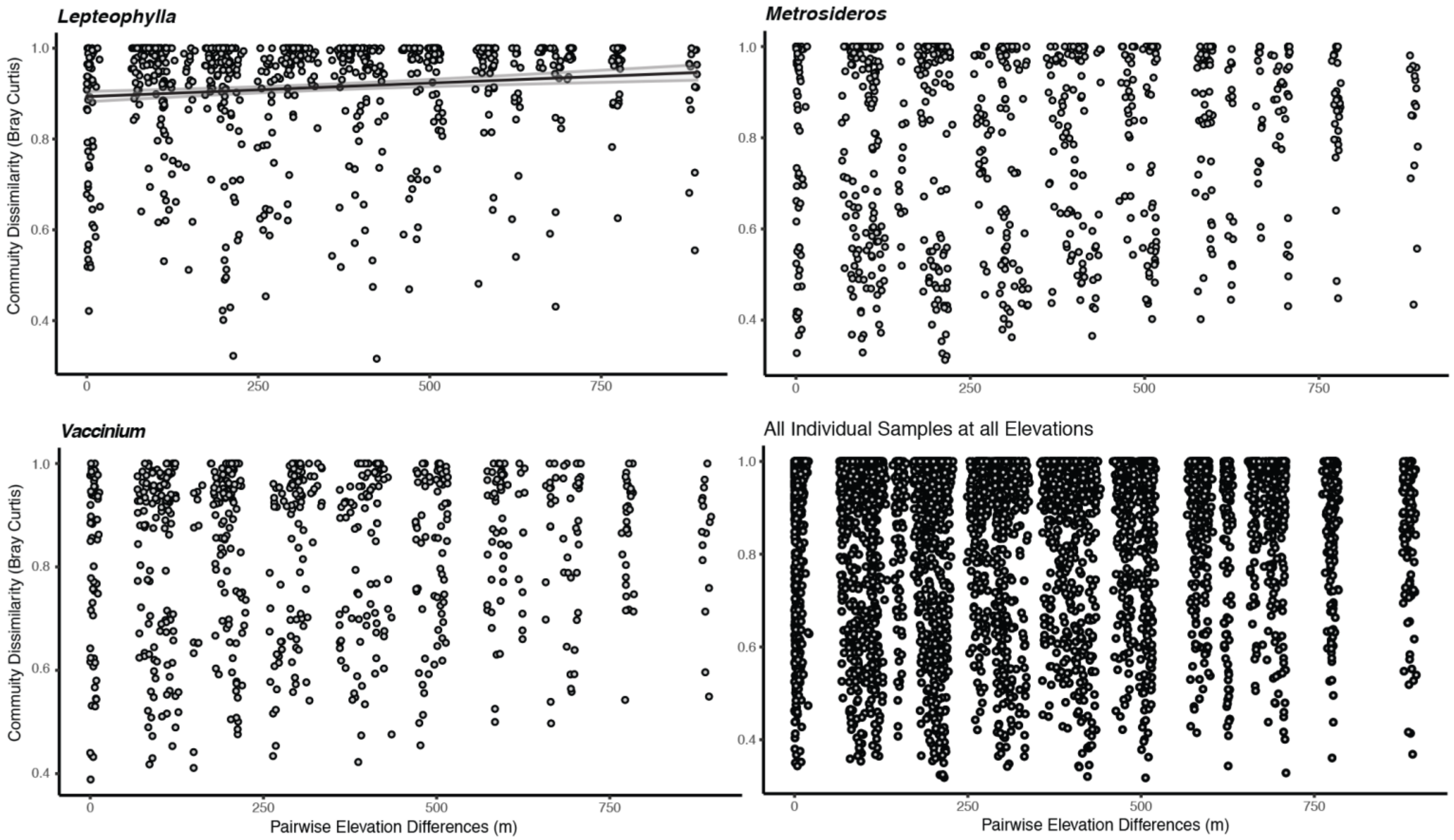


**Figure S2.** Mantel correlation plots shows community dissimilarity as a function of pair-wise elevation differences between two samples (points). There was no significant relationship for all hosts together (*r* = 0.023; *p* = 0.119), *M. polymorpha* (r = 0.099, p = 0.054), nor *V. reticulatum*; (r = 0.089, p = 0.056). There was a significant, though weak relationship among *L. tameiameiae* endophytes (r = 0.105, p = 0.018).
