## Supplementary material for "Plant-microbe specificity varies as a function of elevation": Figure S3

Supplemental Figure 3:

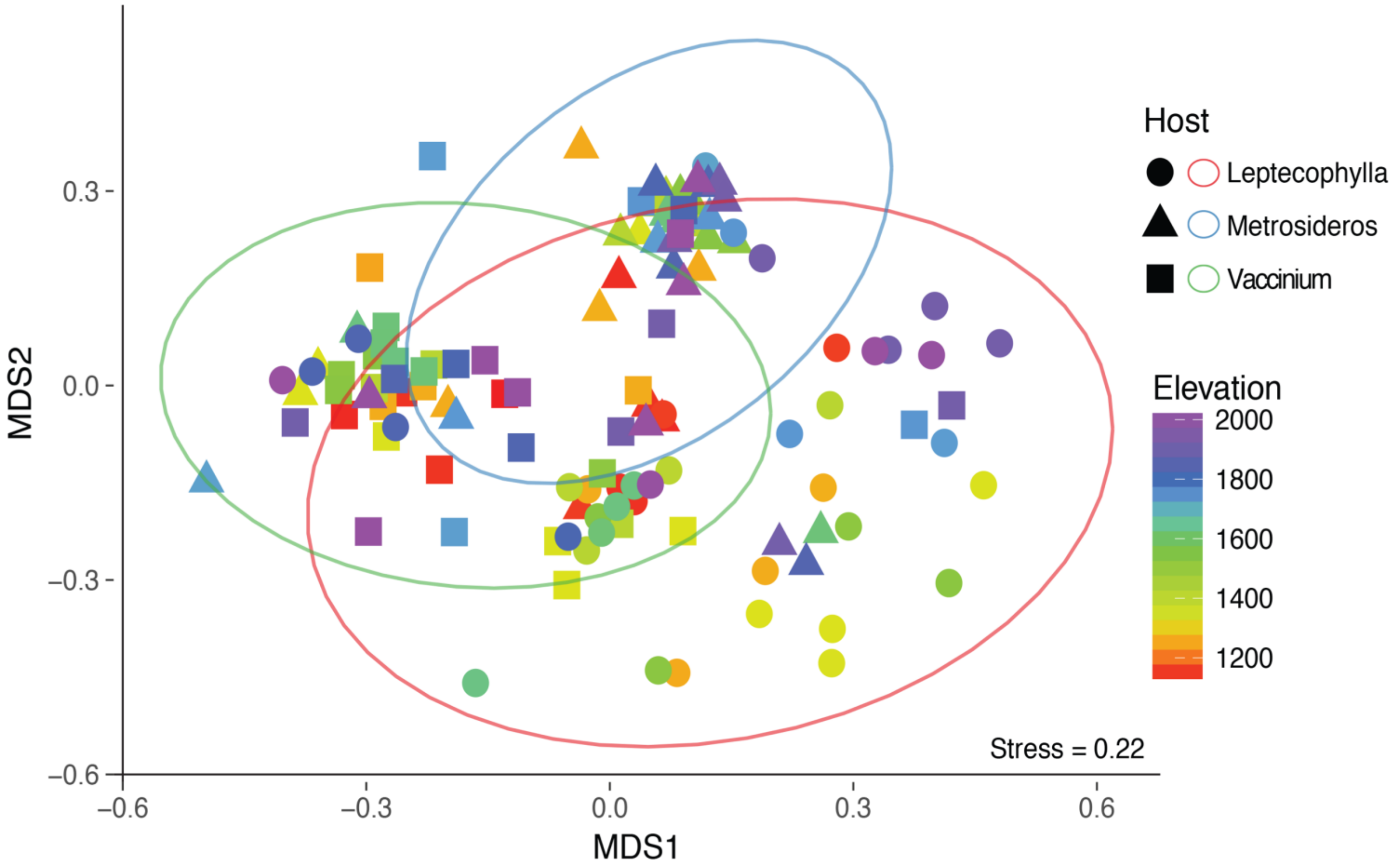


**Figure S3.** NMDS plot for all samples illustrates how hosts weakly structure endophyte communities. Ellipses represent standard error of the mean (95%) for each host. Permutational multivariate analysis of variance was significant for host affects ($R^2$=0.104, p<0.001). Point shapes and colored ellipses correspond to each host species where: squares and red ellipses = *L. tameiameiae*, circles and blue ellipses = *M. polymorpha*, and triangles and green ellipses = *V. retaculatum*.
