## Supplementary material for "Plant-microbe specificity varies as a function of elevation": Table S1

Supplemental Table 1

| **β-diversity Bray-Curtis** | **DF** | **SS** | **MS** | **PsudoF/F** | **R2** | ***p value*** |
|---|---|---|---|---|---|---|
| Host | 2 | 5.003 | 2.502 | 7.388 | 0.104 | 1.00E-04 *** |
| Elevation | 104 | 38.892 | 0.374 | 1.104 | 0.806 | 0.222 |
| Host:Elevation | 7 | 2.678 | 0.383 | 1.13 | 0.055 | 0.228 |
| Residuals | 5 | 1.693 | 0.339 | | 0.035 | |
| Total | 118 | 48.266 | | | 1.000 | |

**Table S1.** Variation in foliar fungal endophyte community composition (beta (β) diversity) among all samples and elevation as determined by permutational multivariate analysis of variance using distance matrices (PERMANOVA). Values shown indicate degrees of freedom (DF), sum of squares (SS), and mean squares (MS). Asterisks indicate significance at $p < 0.05$.
