## Supplementary material for "Plant-microbe specificity varies as a function of elevation": Table S2

Supplemental Table 2

| Site | Observed | Null Mean | t | df | p |
|---|---|---|---|---|---|
| 1100 | 0.6553632 | 0.3314902 | -878.4 | 9999 | < 2.2e-16 |
| 1200 | 0.670467 | 0.3302034 | -822.16 | 9999 | < 2.2e-16 |
| 1300 | 0.8504671 | 0.6075837 | -538.56 | 9999 | < 2.2e-16 |
| 1400 | 0.8656974 | 0.4047464 | -859.21 | 9999 | < 2.2e-16 |
| 1500 | 0.8978409 | 0.5300961 | -583.48 | 9999 | < 2.2e-16 |
| 1600 | 0.8289121 | 0.3618357 | -762.44 | 9999 | < 2.2e-16 |
| 1700 | 0.6024843 | 0.2457483 | -843.64 | 9999 | < 2.2e-16 |
| 1800 | 0.6363899 | 0.237963 | -838.75 | 9999 | < 2.2e-16 |
| 1900 | 0.7322892 | 0.4715782 | -587.16 | 9999 | < 2.2e-16 |
| 2000 | 0.4973674 | 0.2217031 | -853.91 | 9999 | < 2.2e-16 |

**Table S2.** Results of one-sample t-tests between null and observed network specialization (H2’) within sites along the gradient. Observed H2’ values were calculated by aggregating all conspecific samples within an elevation. Null networks were assembled using the swap.web algorithm and the nullmodel function with 10 000 permutations in the bipartite package (Dorman et al. 2008).
