## Supplementary material for "Plant-microbe specificity varies as a function of elevation": Table S3

Supplemental Table 3

| Site | Host | Observed | Null Mean | t | df | p |
|---|---|---|---|---|---|---|
| 1100 | Leptecophylla | 0.638 | 0.3329305 | -564.48 | 9999 | < 2.2e-16 |
| | Metrosideros | 0.548 | 0.3319841 | -403.79 | 9999 | < 2.2e-16 |
| | Vaccinium | 0.782 | 0.3327252 | -841 | 9999 | < 2.2e-16 |
| 1200 | Leptecophylla | 0.757 | 0.3318329 | -702.48 | 9999 | < 2.2e-16 |
| | Metrosideros | 0.652 | 0.3310178 | -535.35 | 9999 | < 2.2e-16 |
| | Vaccinium | 0.604 | 0.3314849 | -453.86 | 9999 | < 2.2e-16 |
| 1300 | Leptecophylla | 0.968 | 0.6069663 | -440.42 | 9999 | < 2.2e-16 |
| | Metrosideros | 0.836 | 0.6077871 | -277.13 | 9999 | < 2.2e-16 |
| | Vaccinium | 0.748 | 0.6094729 | -167.75 | 9999 | < 2.2e-16 |
| 1400 | Leptecophylla | 0.855 | 0.4059882 | -484.94 | 9999 | < 2.2e-16 |
| | Metrosideros | 0.967 | 0.4056056 | -609.08 | 9999 | < 2.2e-16 |
| | Vaccinium | 0.772 | 0.4067242 | -395.91 | 9999 | < 2.2e-16 |
| 1500 | Leptecophylla | 0.862 | 0.532233 | -301.41 | 9999 | < 2.2e-16 |
| | Metrosideros | 0.981 | 0.530428 | -412.48 | 9999 | < 2.2e-16 |
| | Vaccinium | 0.851 | 0.5301789 | -296.26 | 9999 | < 2.2e-16 |
| 1600 | Leptecophylla | 0.966 | 0.3627588 | -661.36 | 9999 | < 2.2e-16 |
| | Metrosideros | 0.715 | 0.3623051 | -387.89 | 9999 | < 2.2e-16 |
| | Vaccinium | 0.807 | 0.3640462 | -479.91 | 9999 | < 2.2e-16 |
| 1700 | Leptecophylla | 0.648 | 0.247192 | -676.45 | 9999 | < 2.2e-16 |
| | Metrosideros | 0.643 | 0.2473202 | -673.77 | 9999 | < 2.2e-16 |
| | Vaccinium | 0.518 | 0.2463585 | -468.01 | 9999 | < 2.2e-16 |
| 1800 | Leptecophylla | 0.712 | 0.2382825 | -746.8 | 9999 | < 2.2e-16 |
| | Metrosideros | 0.814 | 0.2395855 | -894.17 | 9999 | < 2.2e-16 |
| | Vaccinium | 0.386 | 0.2400907 | -223.61 | 9999 | < 2.2e-16 |
| 1900 | Leptecophylla | 0.748 | 0.4718546 | -378.15 | 9999 | < 2.2e-16 |
| | Metrosideros | 0.676 | 0.4728481 | -280.93 | 9999 | < 2.2e-16 |
| | Vaccinium | 0.774 | 0.4717614 | -416.4 | 9999 | < 2.2e-16 |
| 2000 | Leptecophylla | 0.555 | 0.2227885 | -766.87 | 9999 | < 2.2e-16 |
| | Metrosideros | 0.545 | 0.2229063 | -732.6 | 9999 | < 2.2e-16 |
| | Vaccinium | 0.395 | 0.2235372 | -392.25 | 9999 | < 2.2e-16 |

**Table S3.** Results of one-sample t-tests between null and observed host specialization (d’). Results are shown for each host within each site along the gradient. Observed d’ values were calculated by aggregating samples by host at each elevation. Null networks were assembled using the swap.web algorithm and the nullmodel function with 10 000 permutations in the bipartite package (Dorman et al. 2008).
